# Multiplexed Cell-Based Diagnostic Devices for Detection of Renal Biomarkers Using Genetic Circuits

**DOI:** 10.1101/2021.11.14.468540

**Authors:** Sıla Köse, Recep Erdem Ahan, İlkay Çisil Köksaldı, Muazzez Asburçe Bike Olgaç, Çiğdem Seher Kasapkara, Urartu Özgür Şafak Şeker

**Affiliations:** UNAM−Institute of Materias Science and Nanotechnology, National Nanotechnology Research Center, Bilkent University, 06800 Ankara, Turkey; Dr Sami Ulus Children’s Training and Research Hospital, Ankara, Turkey; Ankara Yildirim Beyazit University, Department of Internal Medicine, Children’s Health and Disease Section, Ankara Turkey

## Abstract

The number of synthetic biology based solutions employed in the medical industry is growing every year. The whole cell biosensors being one of them, have been proven valuable tools for developing low-cost, portable, personalized medicine alternatives to conventional techniques. Based on this concept, we targeted one of the major health problems in the world, Chronic Kidney Disease (CKD). To do so, we developed two novel biosensors for the detection of two important renal biomarkers; urea and uric acid. Using advanced gene expression control strategies we improved the operational range and the response profiles of each biosensor to meet clinical specifications. We further engineered these systems to enable multiplexed detection as well as an AND-logic gate operating system. Finally, we tested the applicability of these systems and optimized their working dynamics inside complex medium human blood serum. This study could help the efforts to transition from labor-intensive and expensive laboratory techniques to widely available, portable, low cost diagnostic options.

## Main

Development of novel, versatile, programmable and autonomous diagnostic devices for obtaining a comprehensive view of patient pathophysiology has been a big vacancy in the field of medical diagnostics. Mainly based on analytical chemistry techniques or antibody-based platforms, conventional diagnostic technologies do not match the expected specificity and sensitivity standards, while being low-cost and dynamic[1]. Since the traditional approaches are expensive, time-consuming, require extensive infrastructure and technical expertise, developing new modes of detection for complex and dynamic biomarkers has drawn the attention of synthetic biologists. One of the biological detection tools emerged in this field is whole cell biosensors (WCBs), where biorecognition elements are engineered in genetic circuits to transduce a measurable output in response to desired target molecules[2, 3]. Over the years various cell-based biosensors have been developed for diagnosis of diseases such as inflammation[4, 5], cancer[6, 7], detection of micronutrients[8–10] and pathological agents[11–15]. However, implementing biosensors for diagnostics of complex and interlinked diseases has been especially challenging due to their ambigious nature.

Chronic kidney disease (CKD) emerges from various heterogeneous diseases altering the function and structure of the kidney [16]. CKD is considered a global public health problem, estimated number of people affected reaching 697.5 million worldwide in 2017[17]. Therefore, developing clinically compatible, user-friendly and cost-effective devices targeting CKD hold great potential in this aspect [18].

The end-product of purine metabolism[19], uric acid, is higher in blood serum in humans compared to most other mammals due to the loss of uricase activity by the mutations that happened during the primate evolution[20, 21]. Consequently, humans are more prone to fluctuations in their serum uric acid levels by diet than that in other mammals[22]. Being primarily excreted in kidneys[23] and its relation to complex and bidirectional diseases[24, 25], makes uric acid difficult to identify as a causation or progression factor of CKD. Still, the recent studies have shown that elevated uric acid levels in serum can predict the development of CKD independently. As a result, uric acid is considered an important and potential contributory risk factor for the development and progression of CKD[22].

Urea, a historically significant compound, has been monitored in both urine and in blood for the assessment of kidney function since the mid-19th century[26]. One of the conditions accompanying kidney failure and CKD is the elevated urea levels in the blood. This condition is one of the prominent signs of the impaired kidney clearance function, an irreversible outcome of CKD [27, 28]. While interpretation of blood urea levels have been controversial over the years due to its vulnerability to the environmental factors and dietary intake, Blood Urea Nitrogen (BUN) levels continues to be regularly monitored in patients with kidney related diseases. Furthermore, attractive qualities of urea such as availability at high concentrations and being diffusible, makes measuring clearance rate of urea is a sensitive method for determination of dialysis adequacy for patients with renal failure[29].

Using transcription factor based sensory modules, we have developed gene circuits to detect presence of urea and uric acid. Utilizing synthetic biology inspired methodologies such as promoter engineering, post-transcriptional modifications and protein engineering we have optimized the response curves of each biosensing system. The systems were integrated into a single cell for simultaneous detection. Furthermore, using promoter engineering we also generated an AND-logic gate mimicking system for implementation into synthetic biology inspired biodevices that are capable of complex detection. Finally, for initial validation we have tested our biosensing systems with human serum samples and developed a simple workflow for detection of urea and uric acid using cell-based biosensors.

## Results

Our initial goal was to develop an urea sensitive cellular signal generator through urea dependent gene expression in a synthetic circuit. To do so, we adapted the biological parts found in the urease operon of the uropathogen, *Proteus mirabilis*. Urea is a common environmental trigger for these uropathogens including *P. Mirabilis, Providencia stuartii, Escherichia coli* and *Salmonella species* [30, 31]. The expression of the virulence factor urease in these species is regulated by the urea sensitive transcription factor UreR [32]. One of the members of the AraC family of transcriptional activators, UreR binds to urea and activates the transcription by binding to its cognate promoters located in the intergenic region of the urease operon. Here, UreR is designated as the sensory unit of the biosensor module in the bacterial model *E. coli* by constitutive expression. Signal formation with urea presence was established through the sfGFP expression which is located downstream of the complete intergenic region of *P. mirabilis* with the promoter and regulatory sites.

After establishing a urea responsive biosensing mechanism, a signal amplification modification was done for real life applications. There have been a few studies on the intergenic region of *P. mirabilis* to identify the regulatory regions located as well as the promoters[33, 34]. Since the identified regulatory regions did not fully reflect the behaviour of urea sensitive gene circuits, a genetic insulator was utilized to eliminate post-transcriptional interference of promoter sequences on mRNA. The genetic insulator used in this study, namely, RiboJ is a riboregulator with an autocatalytic function[35]. The genetic insulators have been increasingly used in recently developed gene circuits due to their ability to eliminate context dependence caused by promoter variances[36–38]. Here, RiboJ was placed upstream of RBS structure mitigate the uninteded *cis*-interactions caused by the 5’UTR. The constructed circuit enabled a significant increase in the signal levels upon induction with the same levels of urea. However, as expected, the background signal has also increased with the rise in the translational efficiency of the signalling mRNA.

To increase the dynamic range of the urea biosensor, sensing module UreR was engineered with site-directed mutagenesis. Here based on the functional studies on UreR activity[39], two lysine to alanine mutations were done through site-directed mutagenesis. The first mutation K169A is positioned in the linker region between the dimerization domain and the DNA binding domains of the UreR. The latter mutation K15A on the other hand is in the dimerization domain. The first mutation increased the dynamic range of the urea sensing mechanism to ≈15-fold. Furthermore, the addition of the second mutation resulted in more than 25-fold change. While not certain, the docking studies on UreR-urea with the mutated versions suggest that they might facilitate flexibility of UreR in the unbound state reducing the spontaneous UreR dimerization rate hence lowering the background (data not shown).

Well-known for its withstand to ionizing radiation and many other DNA damaging agents [33, 34], *Deinococcus radiodurans* has an uricase operon that is regulated by the hypothetical uricase regulator (HucR) transcription factor [35]. HucR is a member of MarR family of transcription regulators that generally bind to their cognate promoter/operators as homodimers and repress transcription initiation. Their interactions get negatively affected by the presence of the phenolic ligands which in this case is uric acid [35, 36]. Utilizing this logic, HucR was constitutively expressed in the host with its cognate operator site located in between −35 and −10 regions of its cognate promoter. This minimal promoter was designed to regulate sfGFP expression in the host cell, therefore, uric acid binding to HucR was expected to result in GFP expression regulation. Since the expected induction signal levels were not met, intuitively, processing and sensory units of the biosensing circuit were cloned into a high copy vector. While the HucR transcription was seemingly repressing transcription leading to a low background when the uric acid was absent in the system, the relative induction signal was too low assess the current system as a suitable uric acid biosensor. To solve this problem, without changing the binding dynamics between the TF and its cognate TF-binding site, promoter strength was increased by swapping the native −35 and −10 regions of the pHucR promoter with the highly strong bacteriophage lambda promoter, pL[40]. The results significantly differed compared to previous versions. Host cells carrying uric acid circuit with engineered synthetic promoter have shown extremely low background while response curve has shown a strong digital response to presence of uric acid concentration at μM concentrations.

Fluorescence and the optical density measurements were taken at the 8th hour of induction with urea. Experiments had three replicates and the normalized data was fitted to one site specific binding graph with Hill Slope (GraphPad Prism 8.3).

One of the aims of this study was to apply Boolean logic on medically relevant biomarkers to show robustness of our system. A logic gate mimicking system can help diagnose kidney related disorders more specifically, considering the rise in the levels of biomarkers could also be related to other syndromes[41]. To do so, initially orthogonality of the circuits were tested (Supplementary figure 3). After observing that there is only negligible cross reactivity between urea and uric acid sensor components, a multiplexed system with an AND logic operation capability was developed. For this purpose, in addition to bringing the elements of urea and uric acid sensing mechanism inside the single cell, an AND logic gate where only presence of the both of the inducers activating the processing module was needed. In the preliminary constructs, HucO binding region or double HucO binding region was added downstream of pureD promoter However, trials have shown that the system’s activation was not dependent on uric acid presence (data not shown). Based on uric acid sensing HucR transcription factor’s ability to bind and repress transcription when the operator binding site HucO is between −35 and −10 regions[42], one of the identified promoters in the intergenic region was modified to carry a HucO region between its −35 and −10 regions. This design mimicked AND logic successfully in which it was dependent on urea presence since the promoters UreR on the intergenic region do not show activity without activated UreR homodimer binding.

In the next step, WBCs were multiplexed inside single cells for parallel detection of the renal biomarkers. Since there were not any undesirable interactivity between the circuit components, the sensor elements were cloned into the same cell. The high copy plasmid, pZE, was designed to carry urea biosensor processing unit, uric acid processing unit and uric acid sensory unit. On the low copy plasmid pZS, the urea sensory unit was positioned with uric acid transporter gene. In this dual reporter system, the single analyte circuits had the same systematic design, only difference being the second reporter mScarlet-I gene placement downstream of the uric acid sensing promoter. To increase the urea sensors response levels same rational design strategy was applied to dual reporter biosensor and the insulator RiboJ was cloned downstream of urea sensing promoter.

In the last step of this study, real life applicability of the sensors was put onto the test. To do so, blood serum samples of healthy individuals and individuals with high urea and/or uric acid levels were tested with the single analyte responsive whole cell biosensors. The urea and/or uric acid levels were predetermined with traditional spectrophotometric methods (see Materials and Methods). Due the nature of biosensor sensitivity and the difference between attributed risk levels of chosen renal biomarkers, developed multiplexed systems were inherently unsuitable for detection in single cells. Therefore, single analyte biosensors were used in the optimization process.

While, bacterial cells are capable of growth under varying cultivating conditions, complex environments like blood serum can interfere with operating dynamics of bactosensors. One of the limitations that arise from implementation of real-life, complex samples is that the number of unforeseen effects of different analytes or substances found in the environment on the network of metabolic pathways of living biosensors[43]. Furthermore, the passive immune functionality present in human serum, the complement system, poses another challange that is the inhibition of the bacterial growth[8]. To overcome this issue testing conditions were optimized by varying ratios of serum to minimal medium (Figure S4 and S5). Here, as well as growth, the operational dynamics of biosensors were also optimized by testing three different concentrations (in range, on the border and at the risky levels) chosen from the sample pool. After establishing the coherent response signals at three different concentration, robustness of our biosensors in human serum sample pool with various concentrations was shown by testing 25 serum samples from both healthy and sick individuals(Figure 6).

## Discussion

We have developed and optimized cell-based biosensors for the detection of some of the most routinely screened renal biomarkers[46, 47]; urea and uric acid. Cell-based biosensors show great potential when it comes to reliable, inexpensive and fast-screening biomolecular recognition systems to be applied in the clinical industry as well as the food and environmental industries[48–50]. The developed urea biosensor was initially derived from the urease operon from the organism *P. mirabilis* to function in a genetic circuit implemented in the host *E. coli*. Further, optimization methodology on the initial circuit components with promoter modifications and a genetic insulator increased the output signal more than 6-fold while keeping the relative dynamic range of the biosensor unaffected.

While the genetic insulation tool, RiboJ, is routinely used for circumventing the effects of unintended RNA leaders[36, 51], the recent studies also show that the mRNA transcripts isolated by RiboJ show a general increase in the gene expression[52]. Although the dynamics behind the aforementioned rise in the protein abundance was not clear, the increase in the mRNA stability and its effect on the translational processes are the two of major factors considered[52]. Furthermore, with rational engineering on the UreR transcription factor, a steeper relationship between uninduced and induced states was gained, achieving more than 25-fold change.

We next developed a uric acid sensing circuit in the same host by modifying the HucR-hucO system of *D. radiodurans*. The receptor unit has shown to have an exceptional binding affinity to its operator site in its characterization studies[42] and the measured background of the initial constructs were very low. Since HucR is an allosteric transcription factor that functions via steric hindrance[42], this inherent property of the receptor was utilized in the signal amplification strategy. By modulating the strength of the cognate promoter with viral −35 and −10 regions, the low background of the system was preserved while a dynamic range of 73 fold.

Another novel biosensor developed in this study was the AND-logic mimicking system for urea and uric acid. After careful analysis on the working dynamics of both sensory modules with their respective promoters, a novel promoter was developed by cloning the HucR binding site the Intergenic region of the urease operon with. Even though all the optimization strategies were not transferred into this system in terms of signal amplification, the system has shown the characteristics of an AND-Gate showing great potential in future applications of analyte screening with more complex biomedical algorithms.

In the next step, we developed a multiplexed system to enable parallel measurements of urea and uric acid. After making sure the sensory modules have orthogonality, the biosensing modules were gathered inside a single cell. Reporter molecule of uric acid was replaced with the mScarlet-I to eliminate <u>signal interference between the reporter molecules</u>. Our dual reporter system has shown four distinct phases (no analyte, only urea, only uric acid and both analytes) to the naked eye under the UV transilluminator.

Finally, we have shown functionality of our systems inside medical samples to show its real life implementation potential. Here, multiplexed systems were not suitable for testing due to the difference between the dynamic ranges of the developed sensors and the actual analyte concentration in the human serum samples. Therefore, a coherent and significant response to risky levels of the analytes was achieved with single molecule biosensors and correct concentration adjustments. One of the issues faced was achieving bacterial survival in human serum samples which was overcome by manipulating the ratio between the minimal medium and the human serum amount. However this interferes with the real analyte concentrations versus optimized operational range of biosensors, which shows an adverse effect the real time diagnosis capacity of the biosensors. Therefore, further optimization of testing conditions and a tailored dynamic range is necessary for transitioning these biosensors to point of care devices.

One of the previously recognized drawbacks of transcription factor based biomolecular recognition systems is the improper dynamic ranges which stem from the predetermined sensor characteristics of the derived genetic components [53, 54]. Here, our chosen biomarkers suffered from the same limitation, especially considering the target analytes are naturally circulating in high concentrations in the blood even in healthy individuals. The dynamic ranges of the biosensors could be tailored with external regulatory elements such as a high affinity non-signalling receptor. This then could result in accumulation of the analytes without until the upper limit of health range is surpassed [55]. Furthermore, biosensing circuits could be manipulated to emulate a band-pass filter behavior to achieve a more dynamic tunability and lower the noise.

Cellular biosensors are one of the promising tools that can change the delivery of healthcare. Here, their potential in early diagnosis of renal disorders has been explored with single and multiplexed novel biosensors. Still, there are remaining obstacles that arise from using living organisms as workhorses. However, the new generation of cell based biosensors are emerging with advances in the areas such as microfluidics, 3D bioprinting and microarray technologies[56]. Therefore, our renal biomarker biosensors, when integrated in these recently emerged smart cell culture systems, can enable continuous monitoring of biomarkers in real time without any external effort.

## Materials and Methods

### Strains, Media and Plasmid Construction

*E. coli* DH5α PRO cells were used both for cloning and sensor characterization experiments. Cells were incubated in Luria-Bertani (LB) medium (10 g/l tryptone, 10 g/l NaCl, 5 g/l yeast extract) with their corresponding antibiotics at 37 °C and 200 rpm. Overnight (O/n) cultures were prepared from cell stocks that are preserved in %25 glycerol dissolved in LB medium. O/n cultures are incubated in LB medium for 16h at the aforementioned culturing conditions. For sensor characterizations, (O/n) were diluted 1:100 and grown at the same conditions until their OD_600_ was between 0.4 and 0.6. OD_600_ measurements were conducted with a spectrophotometer (GENESYS 10 Bio, Thermo Scientific).

All primers are listed in Supplementary Table 3.UreR, UACT, HucR, pUreD were synthesized by GENEWIZ. To construct pZS mproD UreR plasmid pZS T7-LacO LuxI plasmid was digested with BamHI and XbaI enzymes. pZE pUreR sfGFP plasmid was digested with BamHI and XhoI enzymes. Single mutations on pZS mproD UreR (K169A) and pZS mproD UreR (K15A-K169A) were introduced with appropriate primers listed in Supplementary Table. RiboJ insulator was added with consecutive PCRs to construct pZE pUreD RiboJ sfGFP. pZS mproD UreR plasmid was digested with HindIII and KpnI and UACT was amplified with appropriate Gibson primers to construct pZS mproD UACT plasmid. pZA AmpR backbone was amplified with appropriate primers and synpHucO was added to sfGFP insert with consecutive PCRs to construct pZA synpHucO sfGFP. proD and HucR regions were inserted into pZA synpHucO sfGFP plasmid after amplification with PCR. pET22b(+) synpHucO sfGFP proD HucR was constructed with PCR amplification of pET22b(+) backbone and synpHucO sfGFP proD HucR amplification. pET22b(+) synpHucO v2 sfGFP proD HucR was constructed with PCR amplification of pET22b(+) backbone and synpHucO sfGFP v2 proD HucR amplification. To construct pZS mproD UreR mproD UACT, pZS mproD was digested with NotI enzyme mproD UACT was amplified with PCR as well as a region of rrnB T1 and T7 terminators. To construct pzE proD HucR pAND sfGFP proD HucR region was added to pZE pUreD sfGFP plasmid, HucO region was added between −35 and −10 regions of pUreD with appropriate primers. pZE proD HucR pUreD sfGFP synpHucO mScarlet-I plasmid was constructed with amplification of mScarlet-I from pEB2-mScarlet-I plasmid with synpHucO promoter and inserted into pZE pUreD sfGFP proD HucR plasmid.

### Sensor Characterizations

Overnight grown cells were diluted 1:100 into 10 mL fresh LB medium with their antibiotic/s and grown until an optical density (OD_600_) of 0.4-0.6 (early exponential phase) was reached. Inducers of indicated concentrations were added from fresh 5M urea stock solution and/or 10mM uric acid solution. Cells were grown at the inducing condition (37°C, 200 rpm) for 16 hours. All experiments were carried out in triplicates.

Whole-cell fluorescence was measured after 0,2,4,6,8,12,16 or 24 hours subsequent to induction for time dependent assays. For dynamic range analysis measurements are taken at the 8^th^ hour (maximum signal amplitude). For fluorescence measurements, an aliquot of 400μL was taken from each sample and centrifuged at 14000 rpm for 3 minutes. Supernatant was removed, and cells were resuspended in 400μL 1x PBS (137 mM NaCl, 2.7 mM KCl, 4.3 mM Na_2_HPO_4_, 1.47 mM KH_2_PO_4_, pH 7.4). 200μL of resuspended cells were transferred to 96-well clear flat bottom polystyrene plates (Corning) for fluorescence measurements.

For all fluorescence and OD_600_ measurements, SpectraMax M5 microplate reader (Molecular Devices, California, USA) was used. sfGFP measurements were conducted with 485-nm excitation filter and 538-nm emission filter with 530nm cut-off. mScarlet-I measurements, excitation wavelength of 544 nm and emission wavelength of 612 nm with 570nm cut-off were used.On the acquired data, signal normalization was done with respect to OD_600_.

### Statistical Analysis

Normalization was carried out by subtracting the value of fluorescence and the value of the OD_600_ of blank PBS solution from the fluorescence value and OD_600_ of each sample respectively. Then, the fluorescence value of each sample was divided by its corresponding OD_600_ value. All data is displayed as mean ± standard deviation. Depending on the data groups, one-way analysis of variance (ANOVA) or two-way ANOVA with Dunnett’s/Tukey’s/Sidak’s multiple comparison test were used (GraphPad Prism 8.3). The min-max normalization of time-dependent fluorescence graph on Figure 5 was calculated by using the following formula;

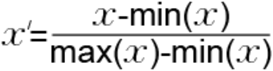

**Figure 1:**
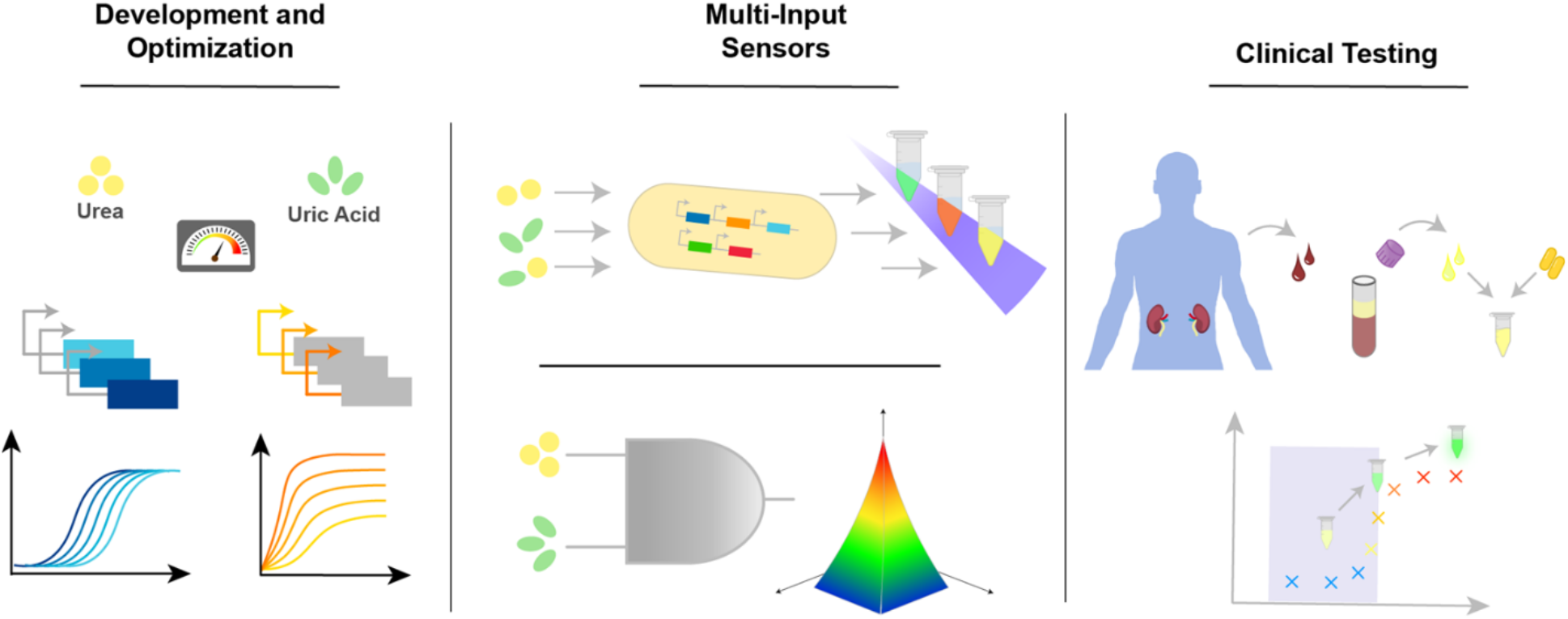
Schematics showing the workflow of cellular biosensor development for medically relevant biomarkers. The cell based sensors are typically generated with a gene circuit carrying a transcription factor (receptor unit), its cognate promoter (processing unit) and a reporter protein (signalling unit). The developed biosensors are designed to respond with a change in the measurable output levels in the presence of the cognate analyte. Both the desired output level and the optimized response curve can be achieved via manipulations on different layers of the synthetic gene circuit as well as the dynamics between these layers. After reaching the ideal sensitivity and operational range from the biosensors, the multi input and logic circuits can be developed according to specific requirements needed for diagnosing the metabolic syndrome of interest. Finally, the biosensors can be tested in the medical medium and testing conditions can be optimized for point-of-care (POC) setups.

**Figure 2.**
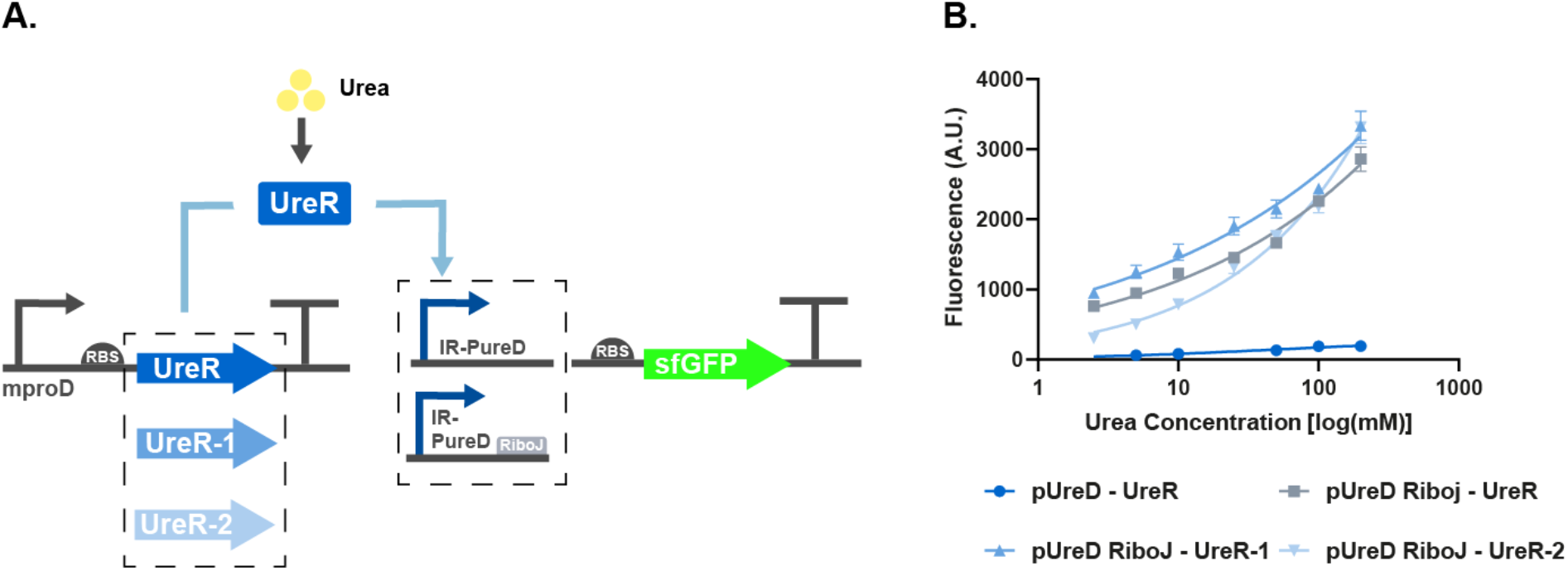
**A**. Schematic representation of the urea sensing mechanism and its optimization steps implemented in the cellular urea biosensor. The receptor unit of the biosensor, UreR, is constitutively expressed inside the cell and activates transcription from its cognate promoter in the presence of urea. Modification steps of the urea biosensor were executed to lower the background expression as well as to amplify the output signal. Via site-directed mutagenesis, different versions of UreR were generated and tested inside the urea sensing circuit. A genetic insulator was deployed to eliminate the unintended 5’UTR interactions with the translation machinery. **B**. The dynamic ranges of the urea biosensors over different concentrations of urea; pUreD-UreR construct shows the detection range of unmodified promoter and the transcription factor. pUreD RiboJ-UreR construct shows the effect of curve RiboJ insulator on the response curve. pUreD RiboJ UreR-1 shows the cumulative effect of the mutation of K169A on UreR gene and the RiboJ insulator on the urea sensor. Finally, UreR-2 shows the double mutated UreR (K169A-K15A) and RiboJ insulator effect on the urea sensor. Fluorescence and the optical density measurements were taken at the 8th hour of induction with urea. Experiments had three replicates and the normalized data was fitted to one site specific binding graph with Hill Slope (GraphPad Prism 8.3).

**Figure 3.**
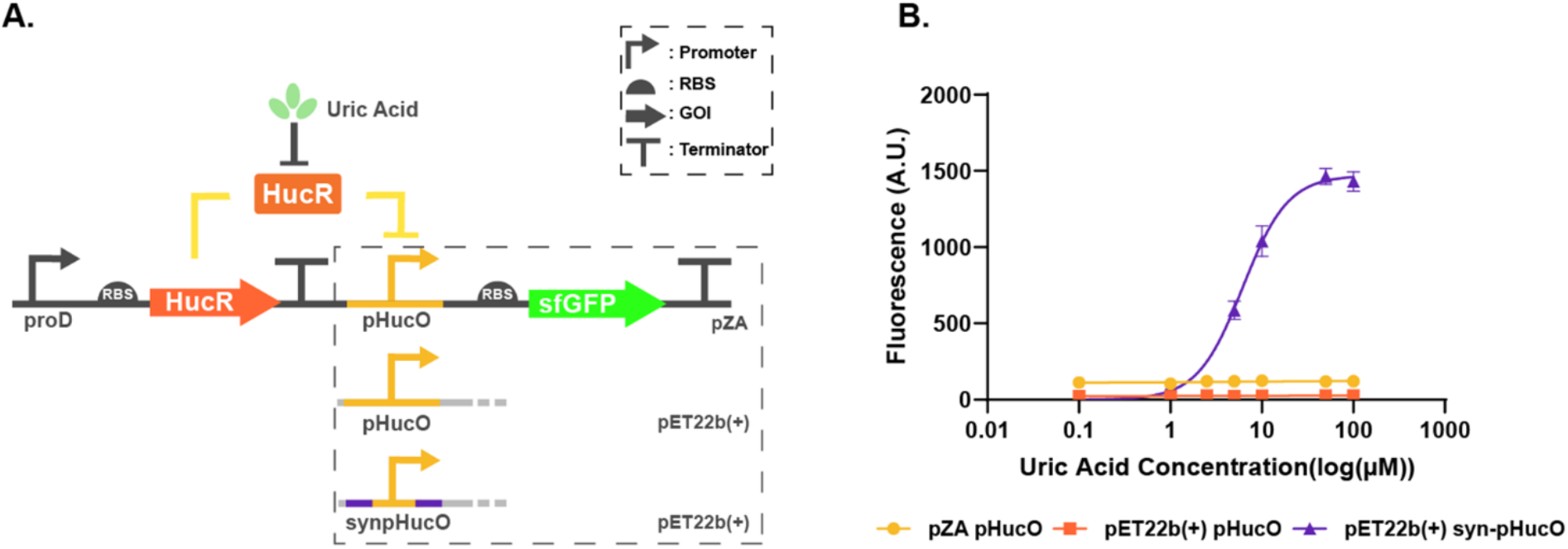
**A**. Depiction of the uric acid sensing mechanism and its optimization steps implemented in the cellular biosensor. The receptor unit of the biosensor HucR is constitutively expressed inside the cell and represses transcription from its cognate promoter in the absence of uric acid. To amplify the signal, the uric acid promoter was designed as a minimal promoter composed of only −35 −10 regions and the HucO binding site of the natural promoter (pZA pHucO). Secondly, sensing and processing units were moved to a high copy plasmid (pET22b(+) pHucO). Finally, the promoter was further engineered with swapping the natural −35 and −10 regions of the synthetic promoter with −35 and −10 regions of the viral strong promoter, pL (pET22b(+) syn-pHucO) **B**. Dynamic Range experiments showing response curves of each uric acid sensing circuit construct; pZA pHucO construct showing minimal uric acid promoter carrying −35 and −10 regions of the natural promoter. pET22b(+) pHucO construct shows the effect of carrying processing and signaling units to a high copy plasmid. Finally, pET22b(+) syn-pHucO construct depicts the cumulative effect of high copy plasmid and engineered synthetic promoter on the overall biosensing efficiency.

**Figure 4.**
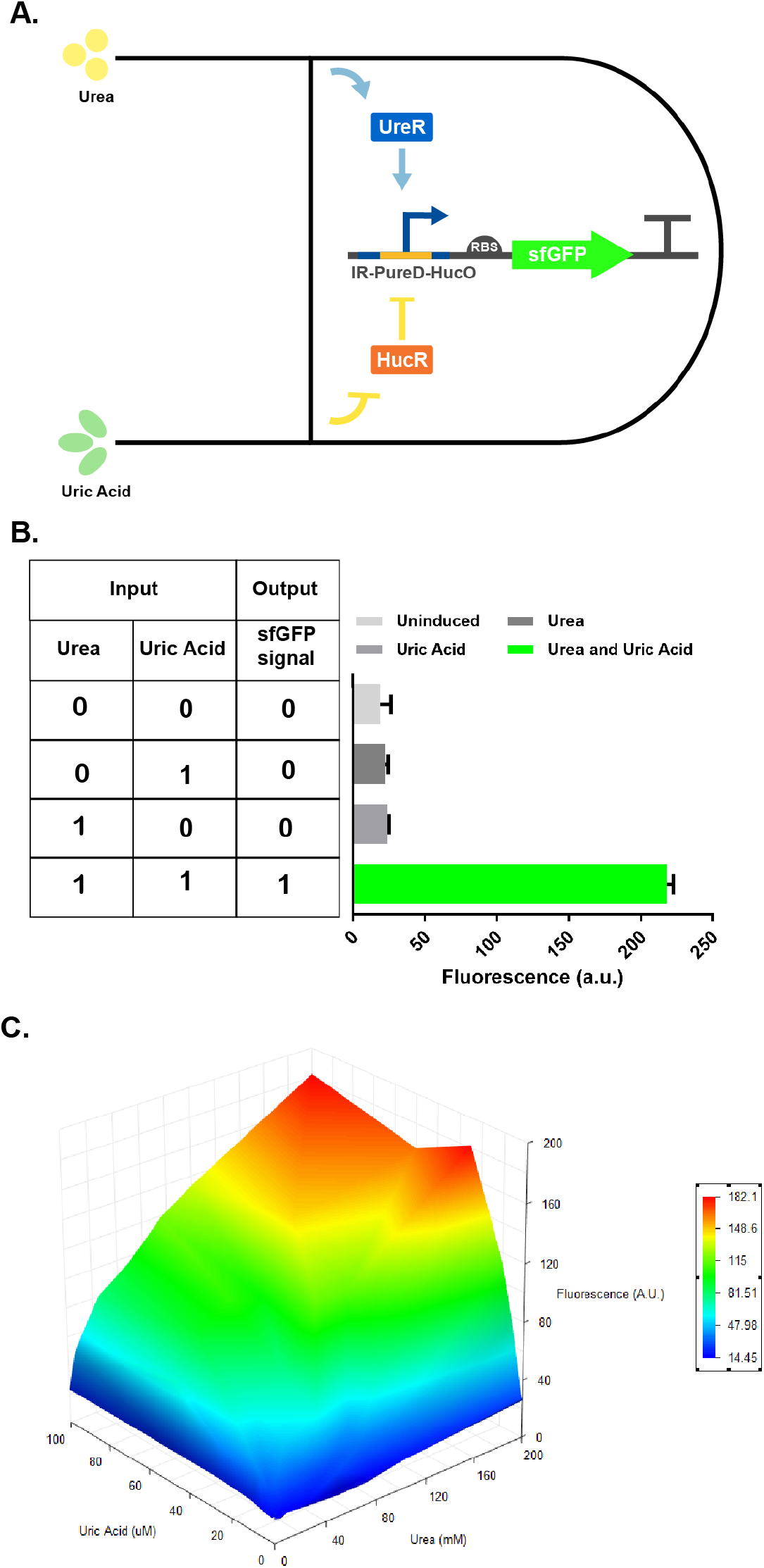
**A**. Schematic representation of the AND logic mimicking biosensor with an AND logic implemented system that is responsive to urea and uric acid. The engineered Intergenic Region from the urease operon which is controlling the sfGFP expression has a HucO operator site in the putative UreD promoter’s −35 and −10 regions. This design prevents transcription from the engineered promoter in the absence of the inputs urea and uric acid by both steric hindrance effect of HucR on the RNAP and the inactive UreR transcriptional enhancer. The presence of either analyte is not is not sufficient to activate transcription however, in the presence of both of the inducers; HucR dissociates from its operator so that the transcription can be initiated by the active UreR. **B**. Fluorescent response graph of AND-Gate biosensor at the 8th hour of induction with the truth table of AND-logic on the left. **C**. Combined dynamic range of AND-Logic Gate Biosensor (0, 5, 10, 25, 50, 100, 200 mM Urea by 0, 0.1, 1, 5, 10, 25, 50, 100 μM Uric Acid). Fluorescence and the optical density measurements were taken at the 8th hour of induction with urea. Experiments had three replicates and the data obtained was normalized (GraphPad Prism8.3).

**Figure 5:**
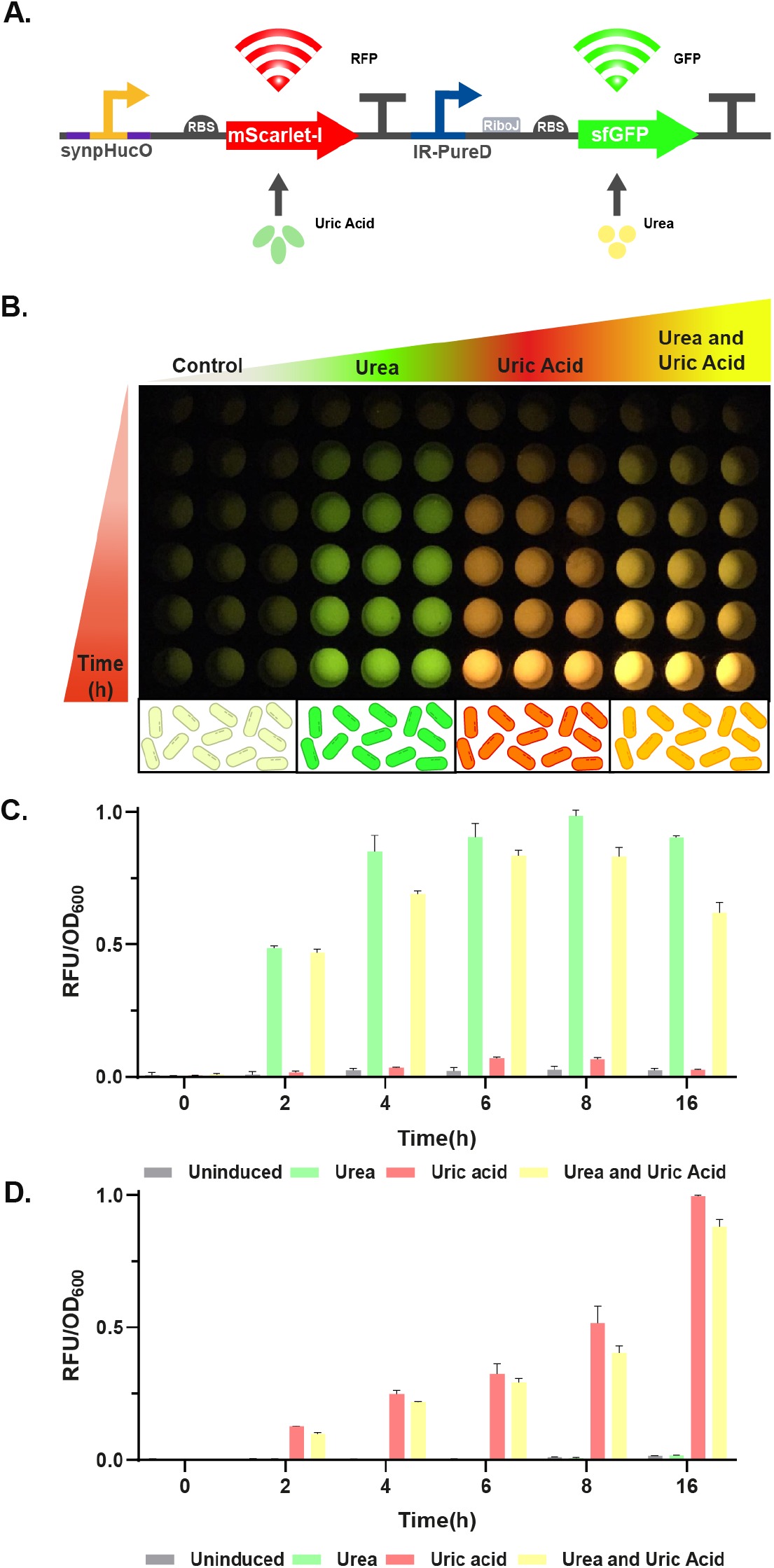
**A**. Schematic depiction of the dual reporter biosensor that is responsive to urea and uric acid. The host cells respond to urea and uric acid by sfGFP expression and mScarlet-I expression, respectively. **B**. Samples in 1xPBS were placed in a 96-well plate before taking the spectrophotometric measurements at different time points. From top to bottom time points depict 0, 2, 4, 6, 8, 16, 24h after induction (Samples were kept in +4°C until the next measurement, picture was taken with a smart phone camera at the last time point). **C**. The time dependent fluorescent graph showing the GFP signal for all cases including uninduced, only urea, only uric acid and both urea and uric acid. **D**. The time dependent fluorescent graph showing the measured RFP signal for all cases including uninduced, only urea, only uric acid and both urea and uric acid. Fluorescence and the optical density measurements were taken at the aforementioned time points with induction cases indicated on the graphs. Experiments had three replicates and the data was normalized (GraphPad Prism8.3) and data fitted to 0-1 scale (Materials and Methods).

**Figure 6:**
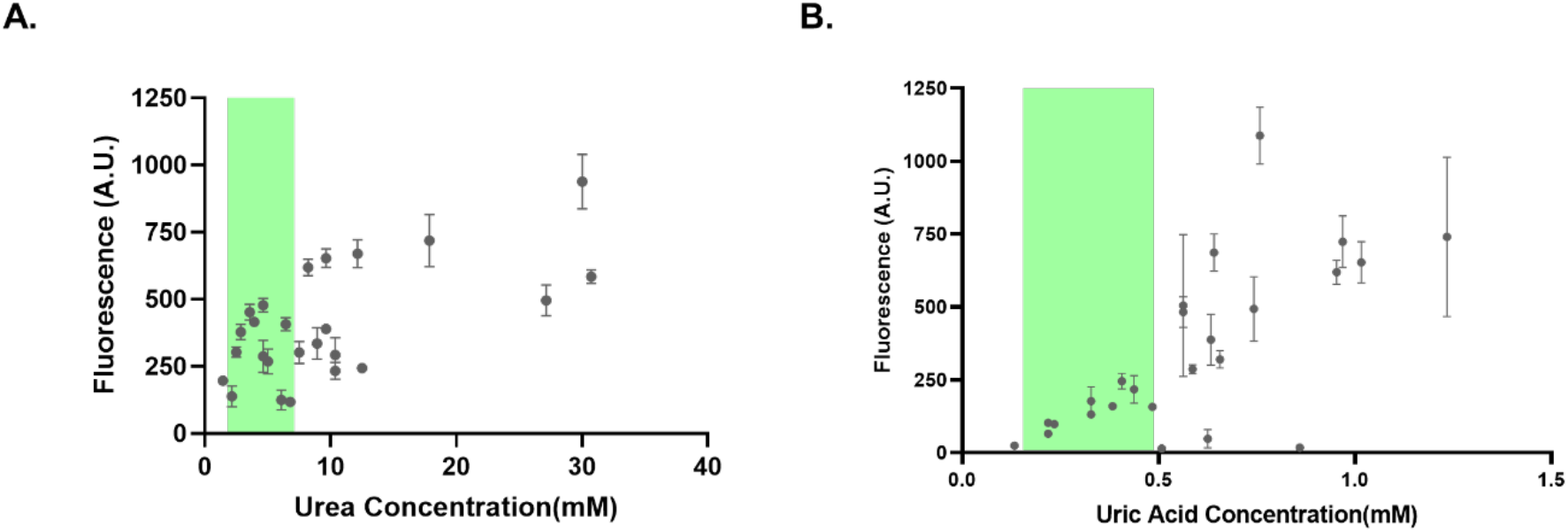
Robustness of bacterial biosensors in human blood serum. **A**. Fluorescent response graph of urea bactosensors inside 5% human serum. The graph shows the measured RFU levels versus the pre-determined urea levels. Green shaded area represents the accepted reference range in this study (1.8-7.1 mM)[44]. **B**. The graph of uric acid bactosensors inside 0.5% human serum. The graph shows the measured RFU levels versus the pre-determined uric acid levels. Green shaded area represents the accepted reference range in this study (155-488μM)[45].

Dynamic range experiments were plotted using One site-Specific binding with Hill slope (GraphPad Prism 8.3).

### Human Blood Serum Assays

Blood serum samples were collected at Dr. Sami Ulus Maternity and Child Health Treatment and Research Hospital, Division of Pediatric Metabolism, Ankara, Turkey under the supervision of Doç. Dr. Asbürçe Olgaç and Doç. Dr. Çiğdem Seher Kasapkara. After obtaining written consent, blood samples were collected by venipuncture of the antecubital vein. The blood samples were centrifuged at 4000 rpm for 10 minutes, and the serum and plasma samples were separated. Blood Urea Nitrogen (BUN) and uric acid values were measured photometrically with the Beckman Coulter AU 5800 (Mishima Olympus Co. Ltd., Japan; Beckman Coulter Inc., CA, USA) analyzer. Remaining serum samples with identified urea and uric acid levels were stored at −20°C until use. Overnight grown urea or uric acid biosensors were diluted (1:100) into M63 minimal medium with corresponding antibiotics and grown at 37°C, 200 rpm until their OD_600_ reached 0.4-0.6. Samples with 300 μL volume were divided into 1.5 mL eppendorfs. Cells were then centrifuged at 3000 rpm for 10 minutes. Supernatant was removed and the cell pellet was resuspended in appropriate volume of fresh M63 medium and serum samples to have 300 μL final volume. Biosensors were induced with human serum samples that have different levels of urea and uric acid. Urea biosensors were induced with addition of 15 μL serum sample (serum 1:20 diluted) unless otherwise stated. Uric Acid biosensors were induced with addition of 1.5 μL serum sample (serum 1:200 diluted) unless otherwise stated. Biosensors were induced for 16 hours unless stated otherwise. Fluorescence measurements were conducted as described above.

### Human Blood Serum Test Optimization Assays

Overnight grown urea or uric acid biosensors were diluted (1:100) into M63 minimal medium with corresponding antibiotics and grown at 37°C, 200 rpm until their OD_600_ reached 0.4-0.6. Samples with 300 μL volume were divided into 1.5 mL microfuge tubes. Cells were then centrifuged at 3000 rpm for 10 minutes. Supernatant was removed and the cell pellet was resuspended in appropriate volume of fresh M63 medium and varying volumes of serum samples to have 300 μL final volume. At different concentrations of serum blood serum samples from the same three patients were used to eliminate interference of other factors such as varying pH, other analyte levels and purity of blood serum. Biosensors were induced with human serum samples that have three different levels of urea and uric acid; on the reference range, near the border and at the risk levels. Urea biosensors were induced with addition of 15 μL serum sample (serum 1:20 diluted) or 30 μL serum sample (serum 1:10 diluted). Uric Acid biosensors were induced with addition of 1.5 μL serum sample (serum 1:200 diluted) or 1.5 μL serum sample (serum 1:100 diluted). Fluorescence and optical density measurements were carried out as described above.

## Supplementary Information

**Figure S1:**
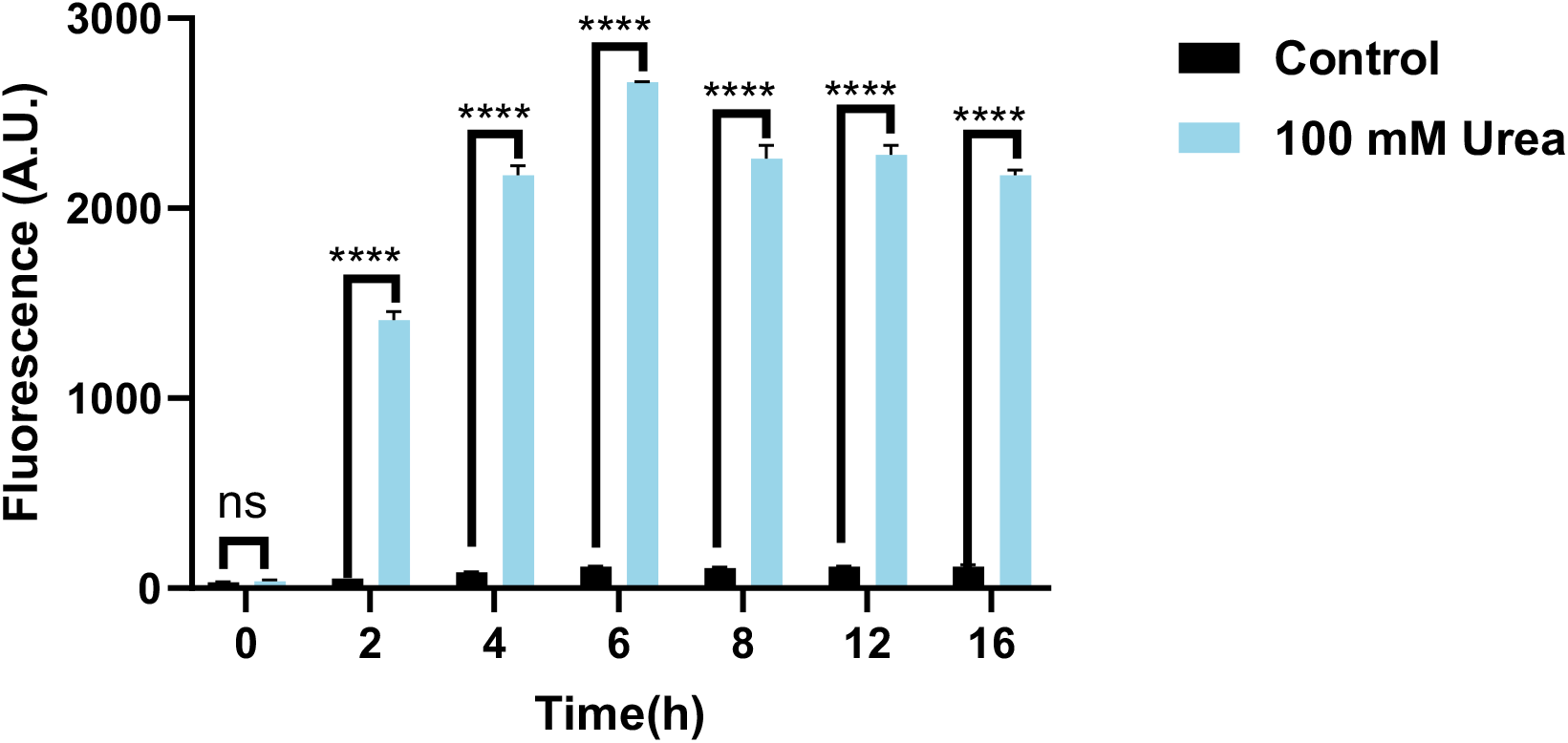
Characterization of the pZS mproD mt UreR (K15A-K169A) urea biosensor with pZE IR pUreD RiboJ sfGFP. After induction with 100 mM urea solution the response signal was measured with a microplate reader at the 0th, 2nd, 4th,6th,8th,12th and the 16th hours. Experiments were conducted with three replicates and the normalized data was analyzed with two-way ANOVA (p ≤ 0.05, p ≤ 0.01, p ≤ 0.001 and p ≤ 0.0001 were shown as “*”, “**”, “***” and “****” respectively).

**Figure S2:**
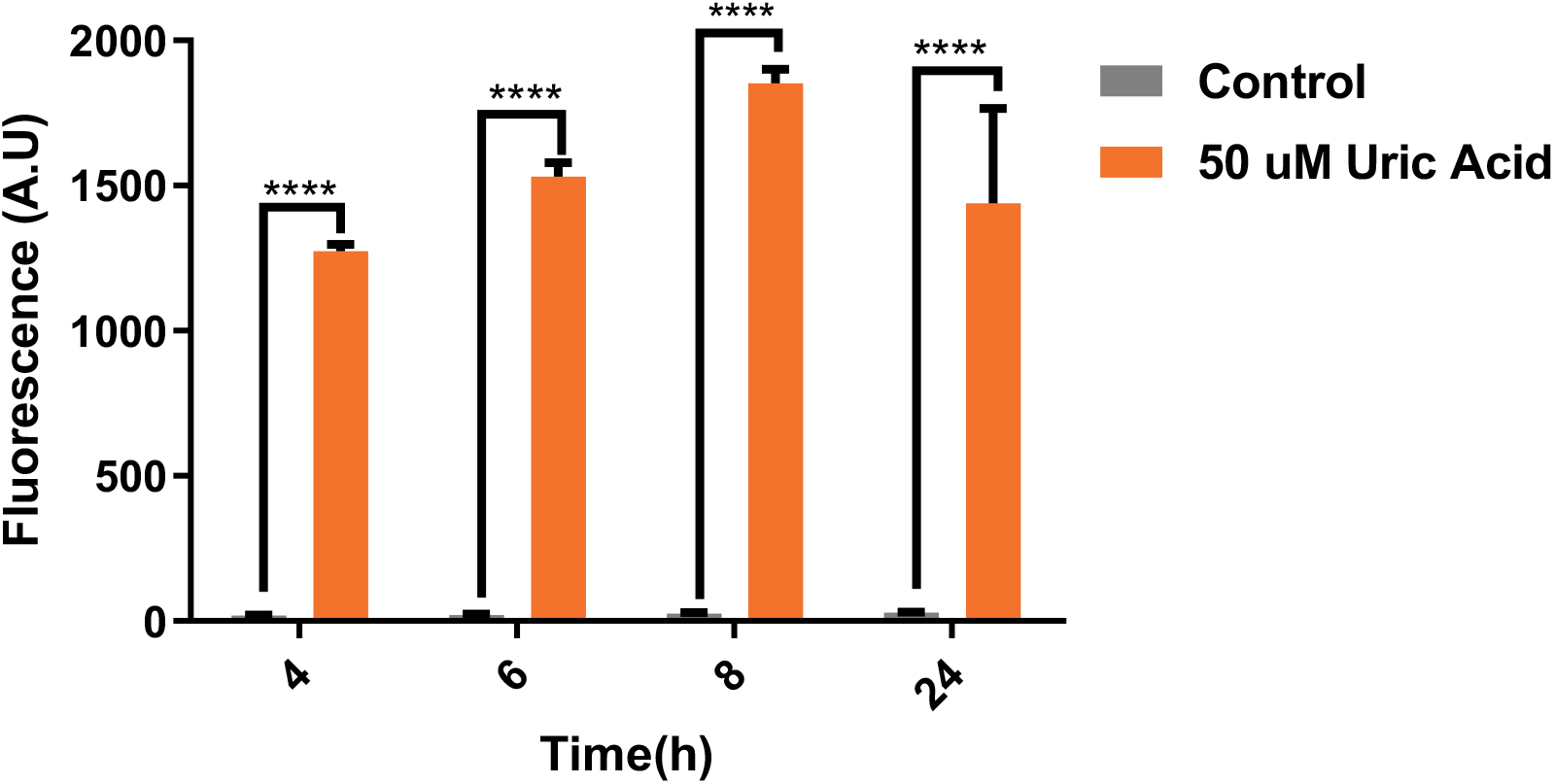
Characterization of the uric acid biosensor. Host cells wereco-transformed with pET-22b(+) synpHucO v2 sfGFP proD HucR and pZS mproD UACT plasmids. After induction with 50 _M uric acid solution the response signal was measured with a microplate reader at the 4th,6th,8th and the 24th hours. Experiments were conducted with three replicates and the normalized data was analyzed with two-way ANOVA (p ≤ 0.05, p ≤ 0.01, p ≤ 0.001 and p ≤ 0.0001 were shown as “*”, “**”, “***” and “****” respectively).

**Figure S3:**
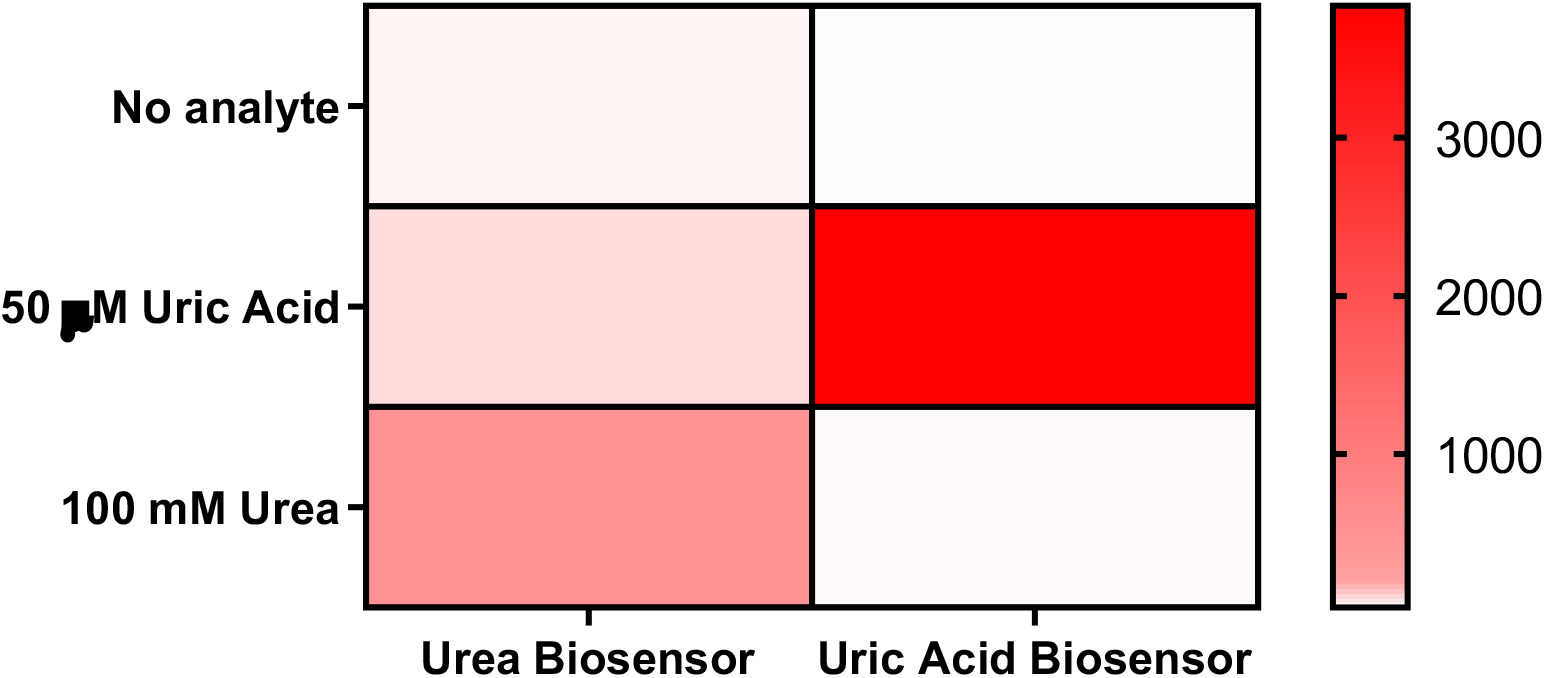
Heat map showing the cross reactivity of individual biosensors to each other’s analyte. To test urea biosensor, UACT gene was cloned downstream of the mproD promoter of the plasmid carrying mproD UreR. The plasmids pZE pUreD sfGFP and pZS mproD UreR mproD UACT were co-transformed into the host cells to permit uric acid flux to the intracellular environment. Co-transformed cells were then induced with 50 μM uric acid or 100 mM urea(left). To test uric acid biosensor, pET-22b(+) synpHucO v2 sfGFP proD HucR and pZS mproD HucR co-transformed into host cells. Co-transformed cells were then induced with 50 μM uric acid and 100 mM urea(right). After induction the response signal from uninduced, uric acid-induced and urea-induced samples was measured with a microplate reader at the 8th hour. Experiments were conducted with three replicates.

**Figure S4:**
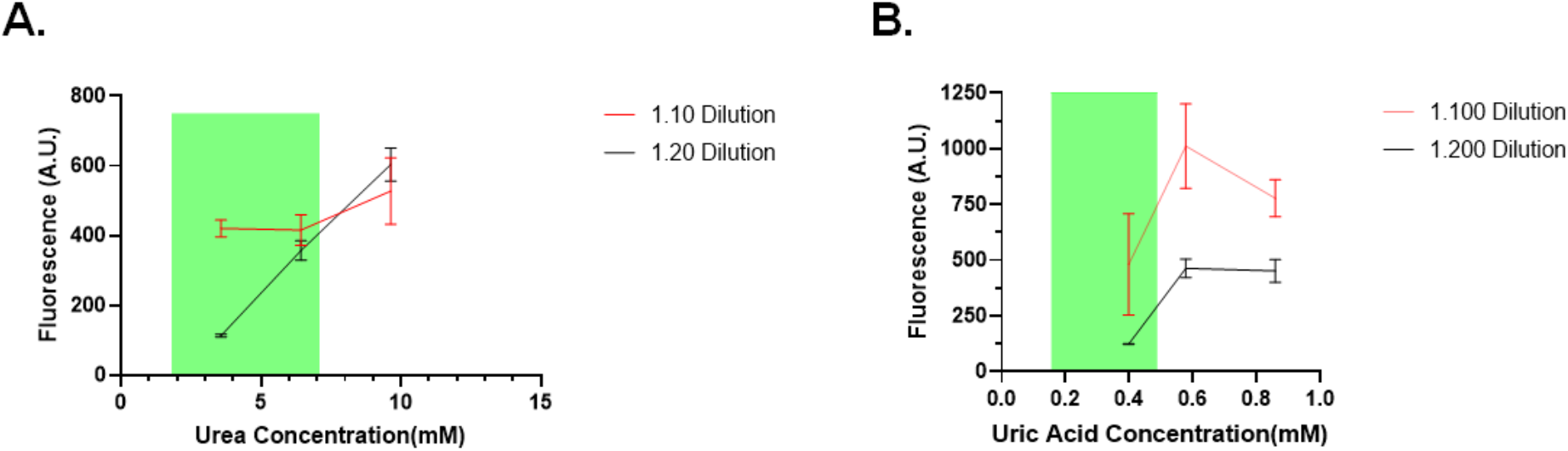
Optimization on testing conditions. **A**. Fluorescent response graph of urea bactosensors inside 5% and 10% human serum. The graph shows the measured RFU levels versus the predetermined uric acid levels. Green shaded area represents the accepted reference range in this study (1.8-7.1 mM)[44]. **B**. Fluorescent response graph of urea bactosensors inside 0.5% and 1% human serum. The graph shows the measured RFU levels versus the predetermined uric acid levels. Green shaded area represents the accepted reference range in this study (155-488μM)[45].

**Supplementary Table 1:** Concentration of urea in the serum samples of patients chosen for the optimization process of testing conditions. Respective concentration changes with every dilution factor are also calculated and depicted.

| Samples<br>(Reference Range:<br>1.8-7.1 mM) | Measured Urea Level<br>(mM) | Urea Concentration<br>in the Sample<br>(Dilution Factor 1.10) | Urea Concentration<br>in the Sample<br>(Dilution Factor 1.20) |
| --- | --- | --- | --- |
| In the Reference Range | 3,57 mM | 0.36 mM | 0.18 mM |
| Near the Risk Border | 6,43 mM | 0.64 mM | 0.32 mM |
| Risky Level | 9,64 mM | 0.96 mM | 0.48 mM |

**Supplementary Table 2:** Concentration of uric acid in the serum samples of patients chosen for the optimization process of testing conditions. Respective concentration changes with every dilution factor are also calculated and depicted.

| Samples<br>(Reference Range:<br>0.155-0.488 mM) | Measured Uric Acid<br>Level (mM) | Uric Acid Concentration<br>in the Sample<br>(Dilution Factor 1.100) | Uric Acid Concentration<br>in the Sample<br>(Dilution Factor 1.200) |
| --- | --- | --- | --- |
| In the Reference Range | 0.398 mM | 0.004 mM | 0.002 mM |
| Near the Risk Border | 0.578 mM | 0.006 mM | 0.003 mM |
| Risky Level | 0.859 mM | 0.009 mM | 0.004 mM |

**Table A.1:**
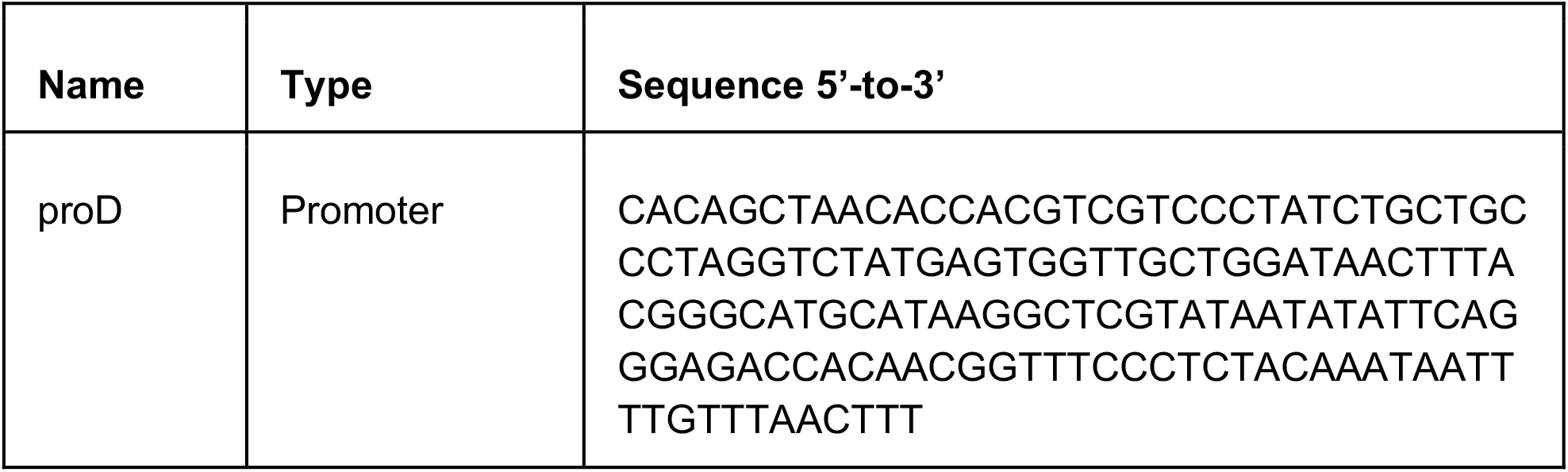
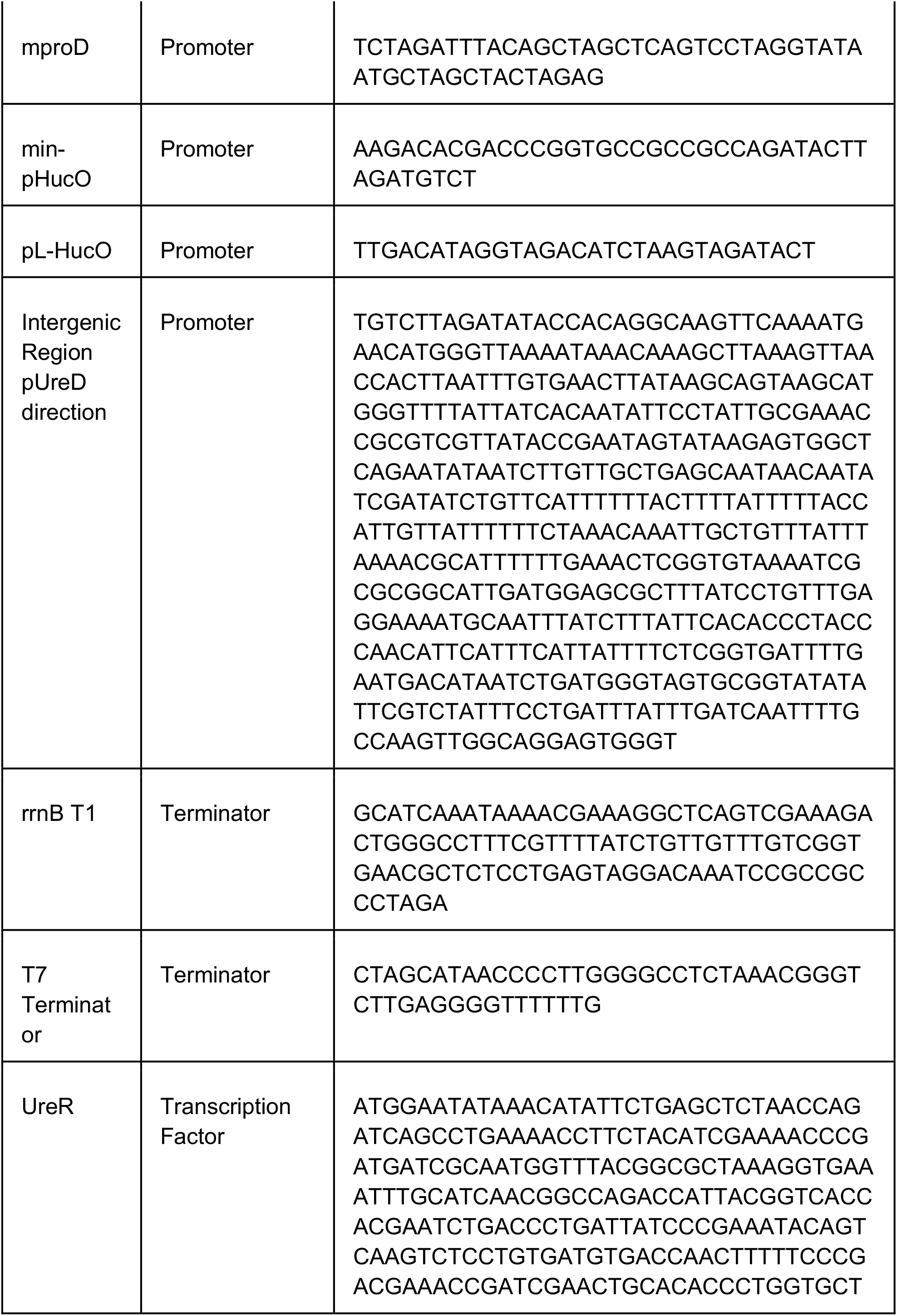
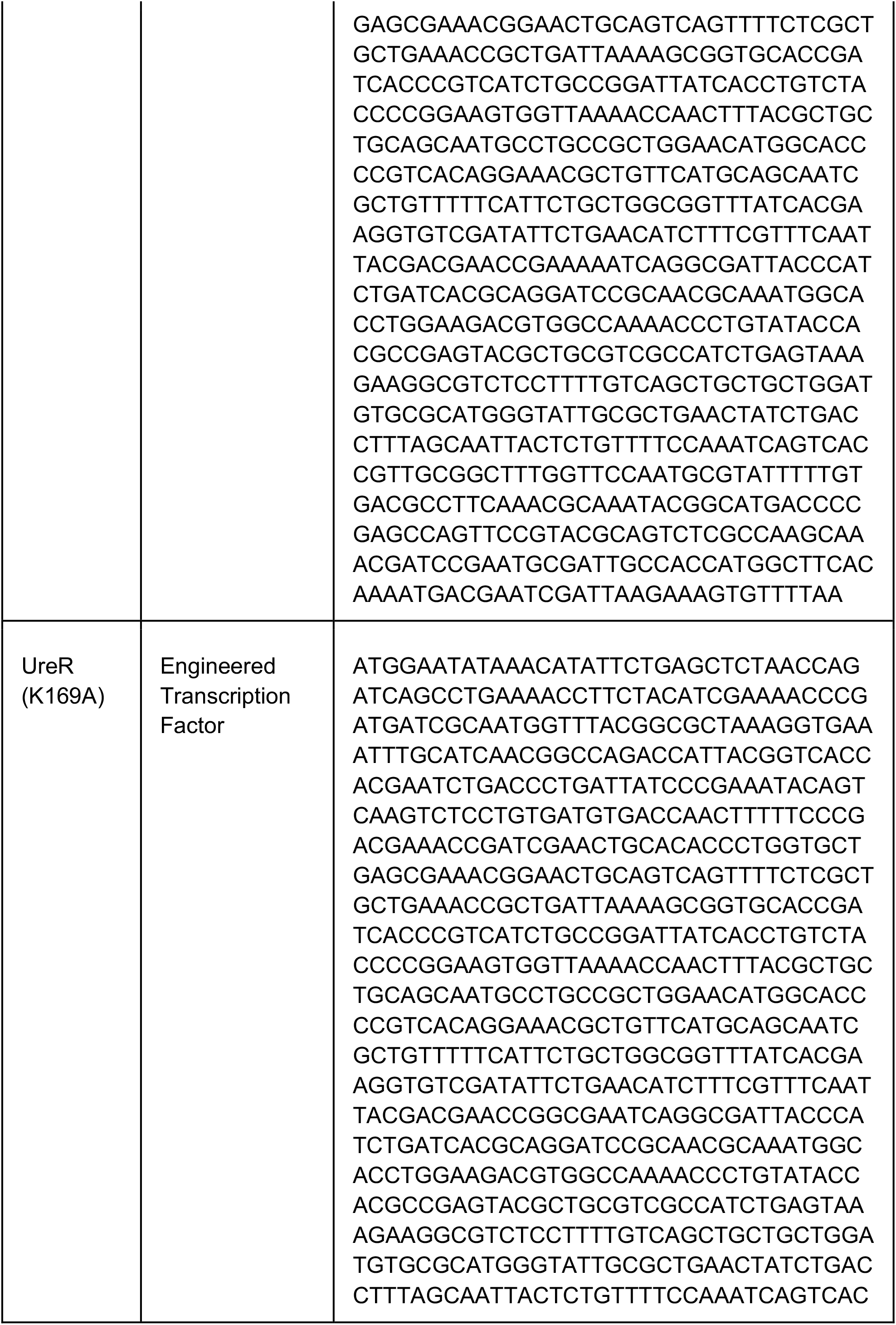
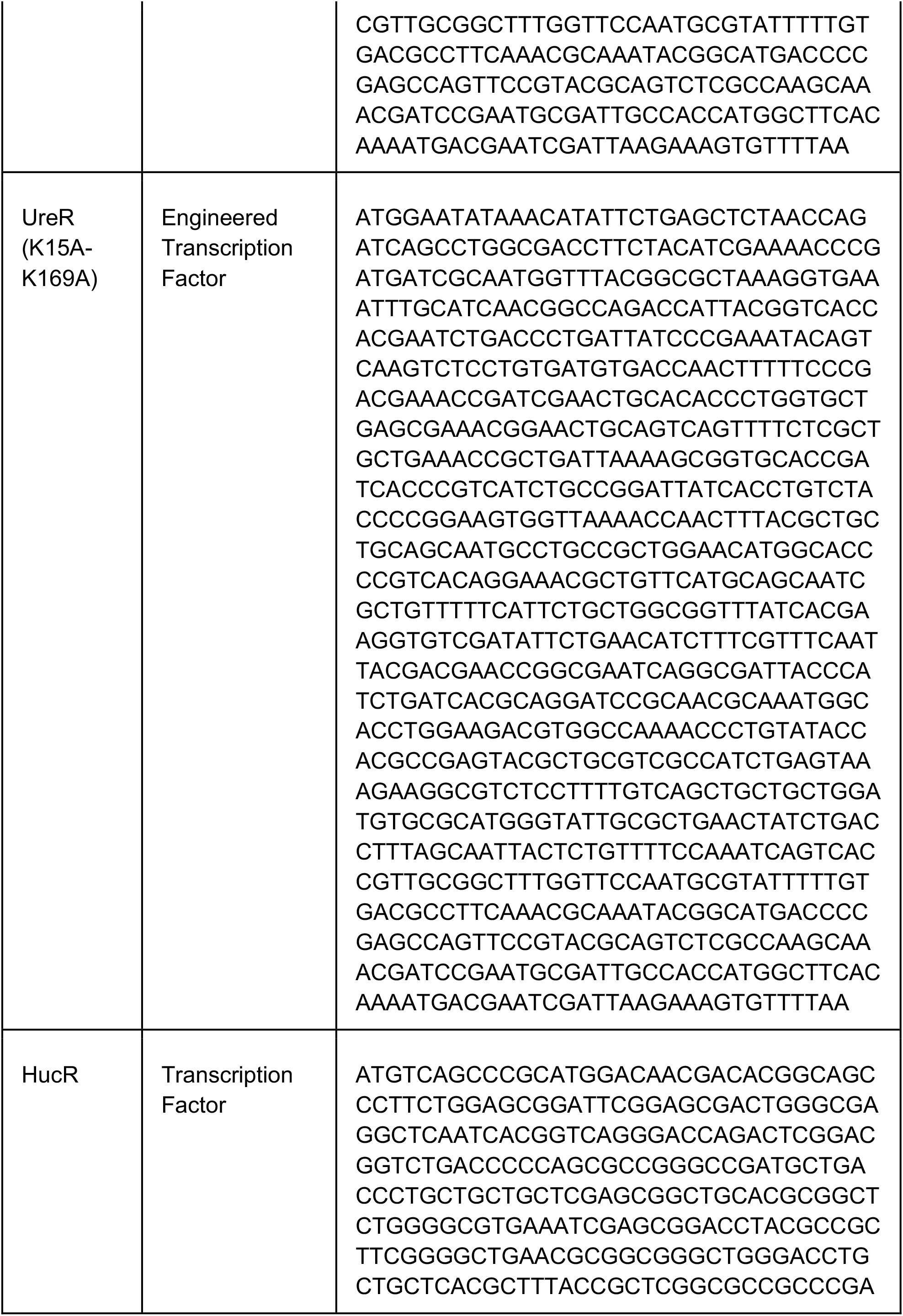
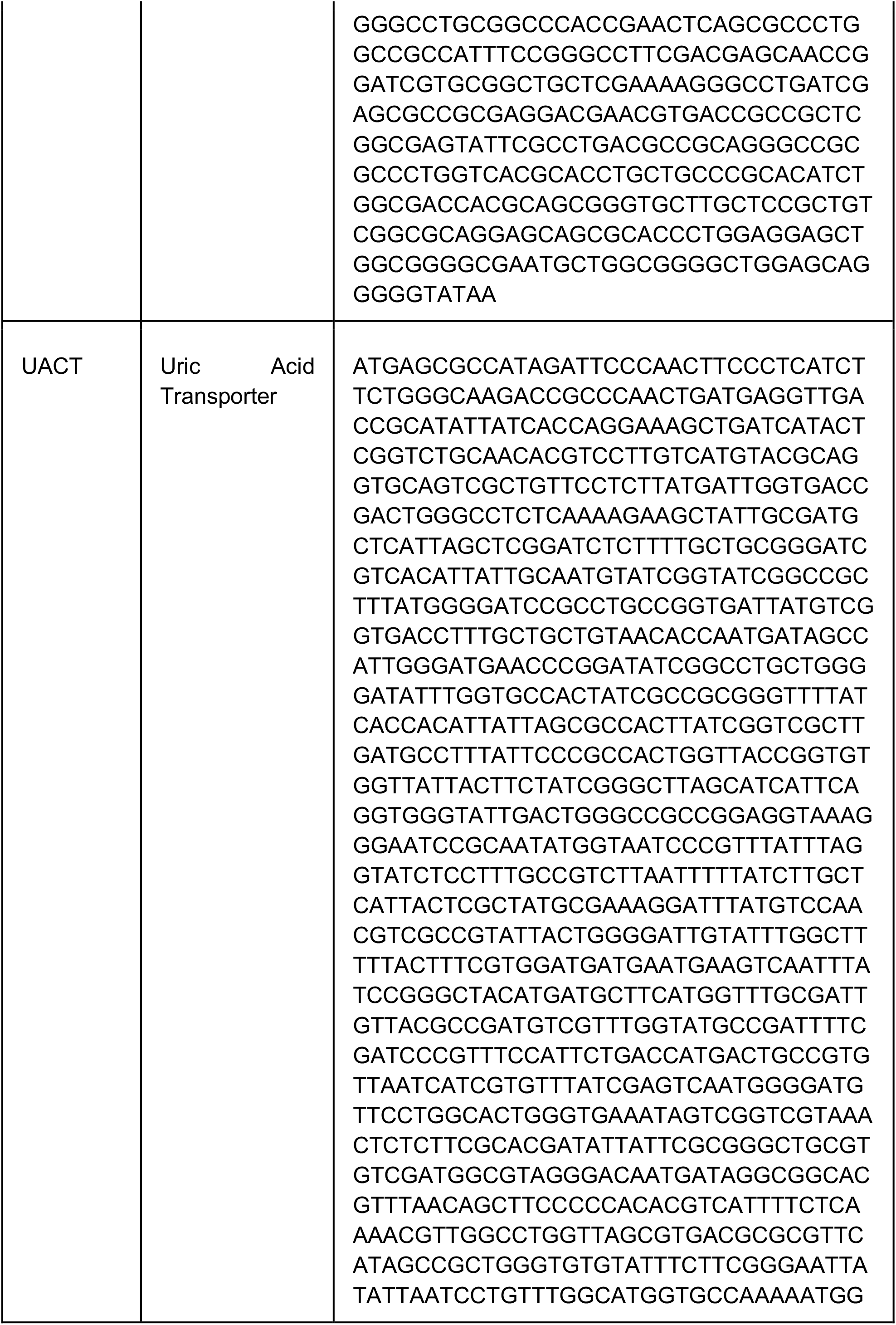
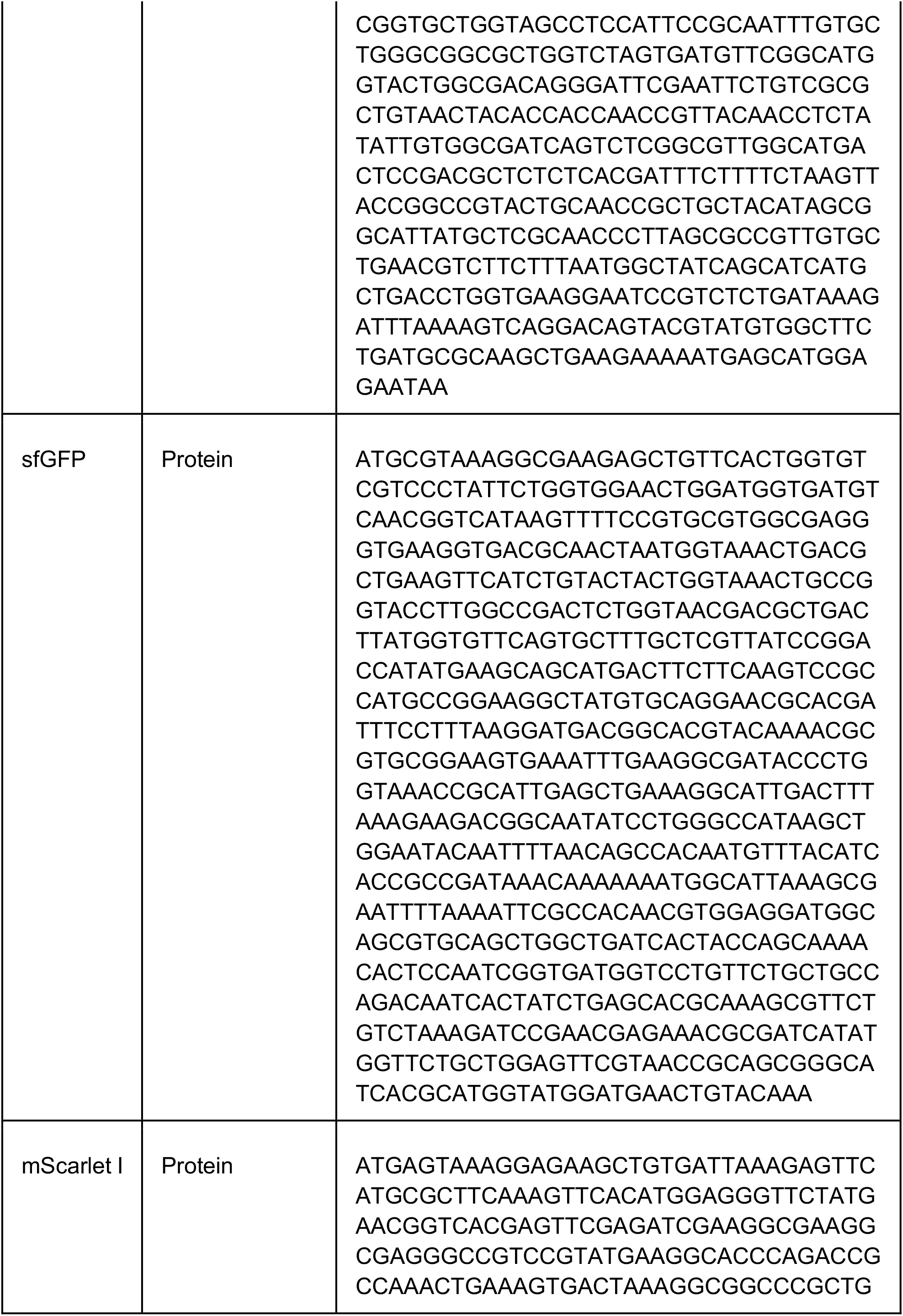
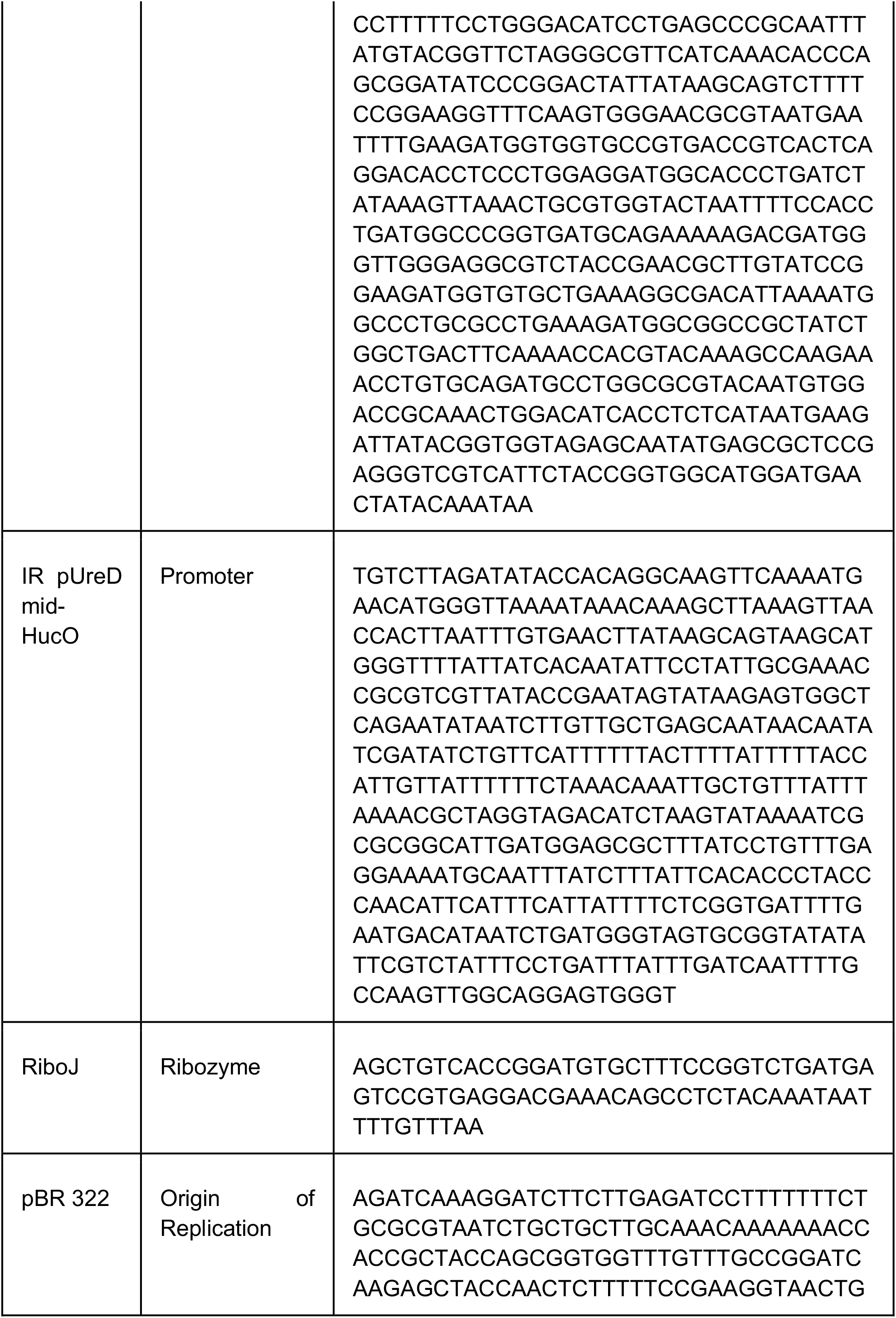
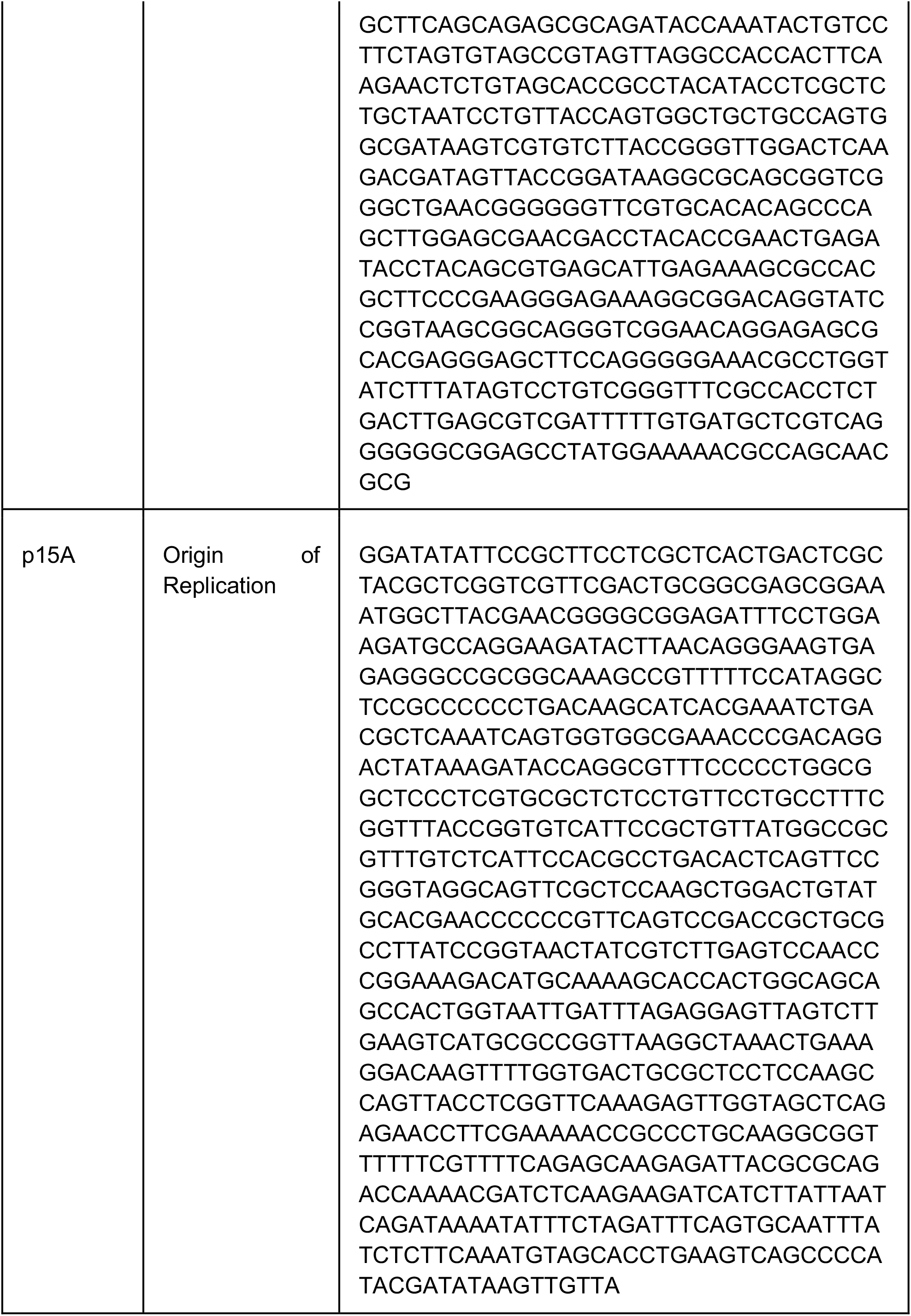
Sequences of genetic parts used in this study.

**Supplementary Table 3:**
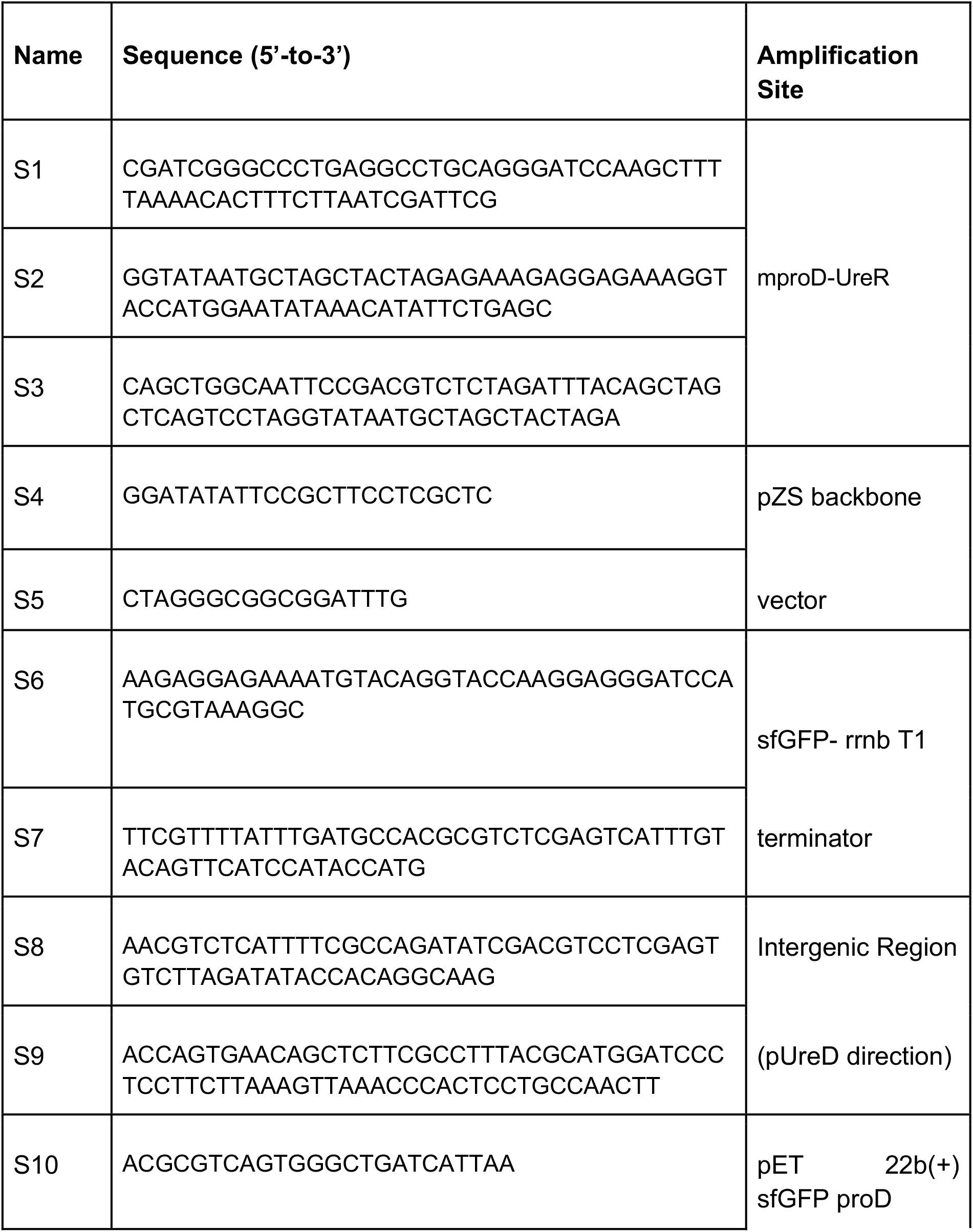
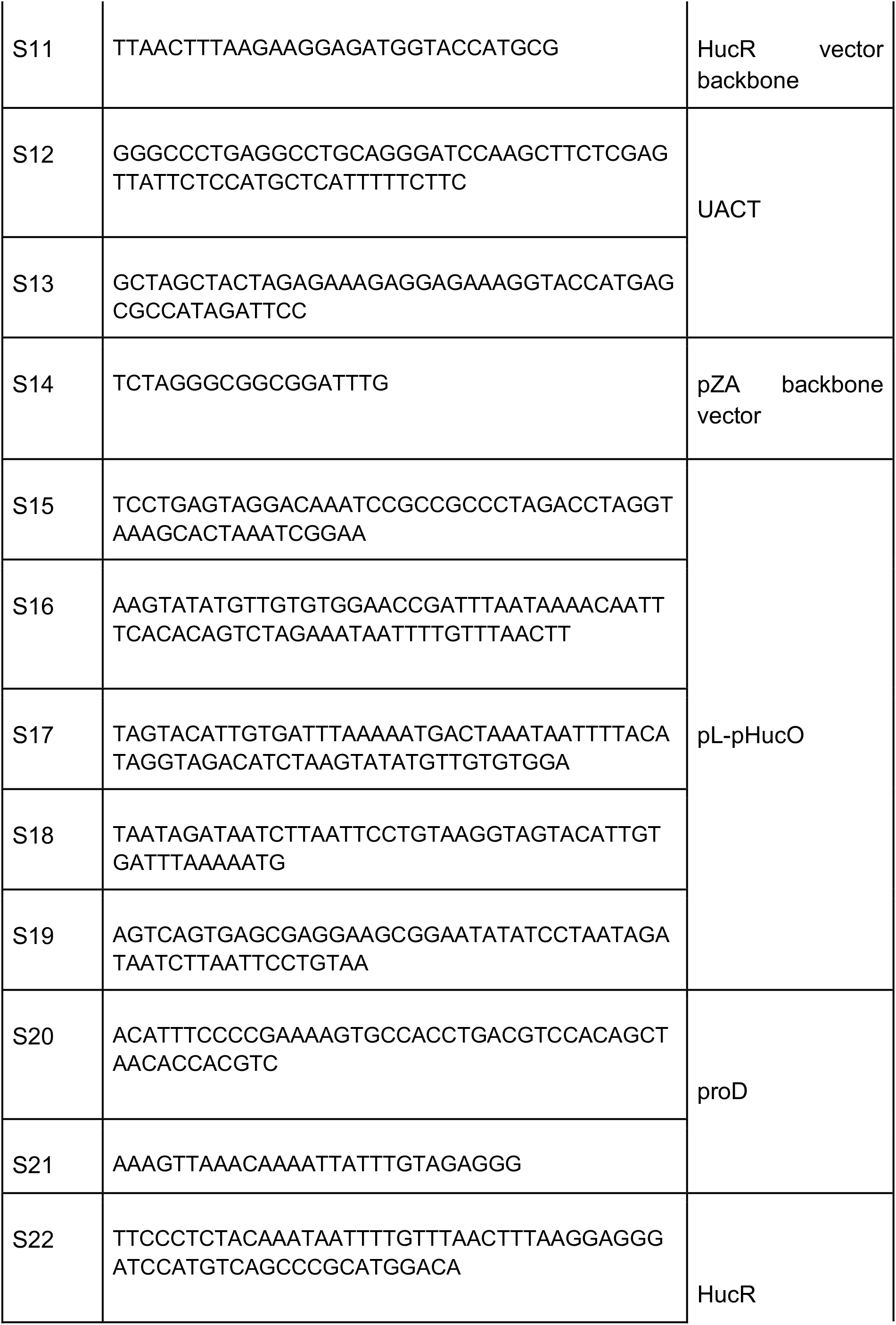
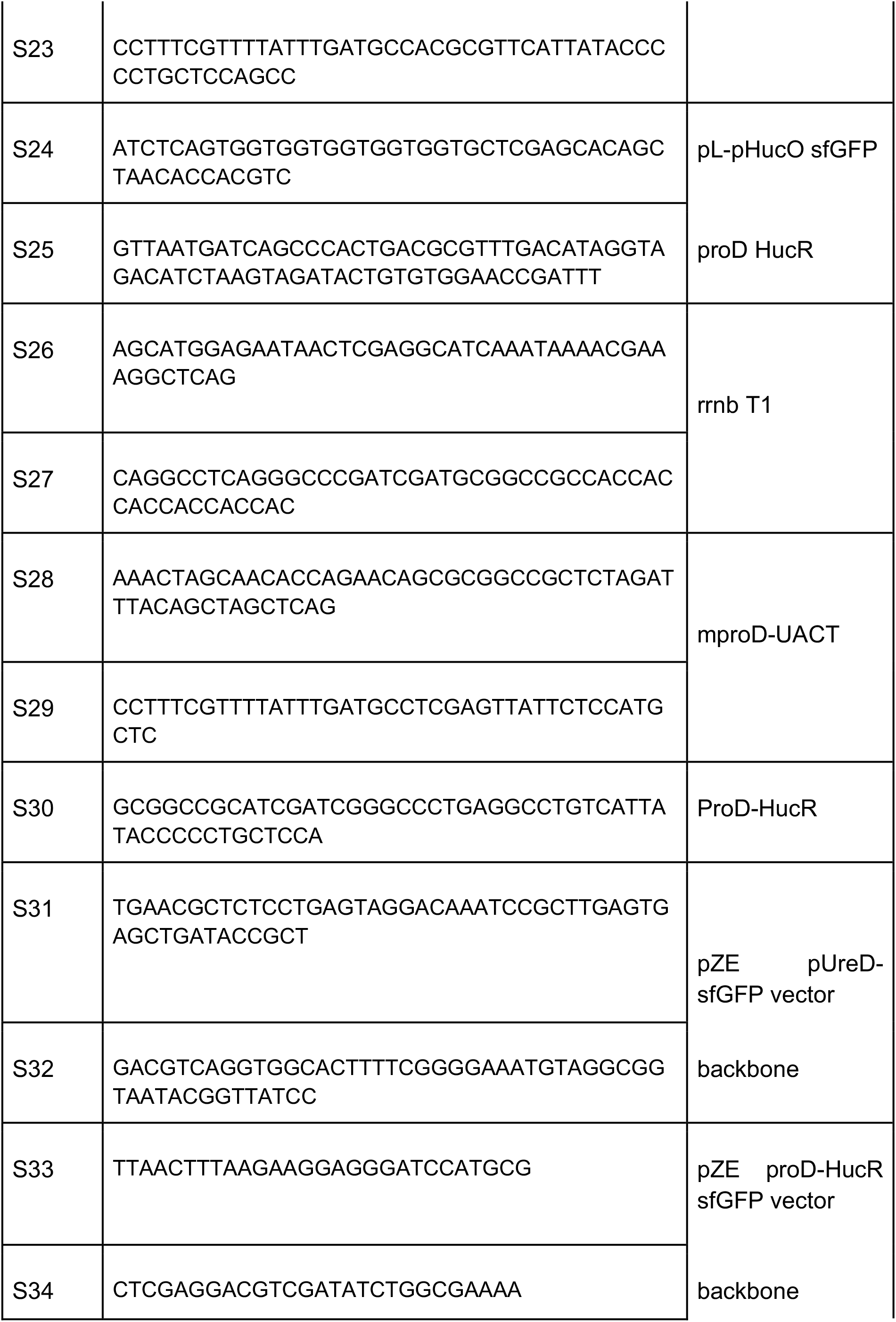
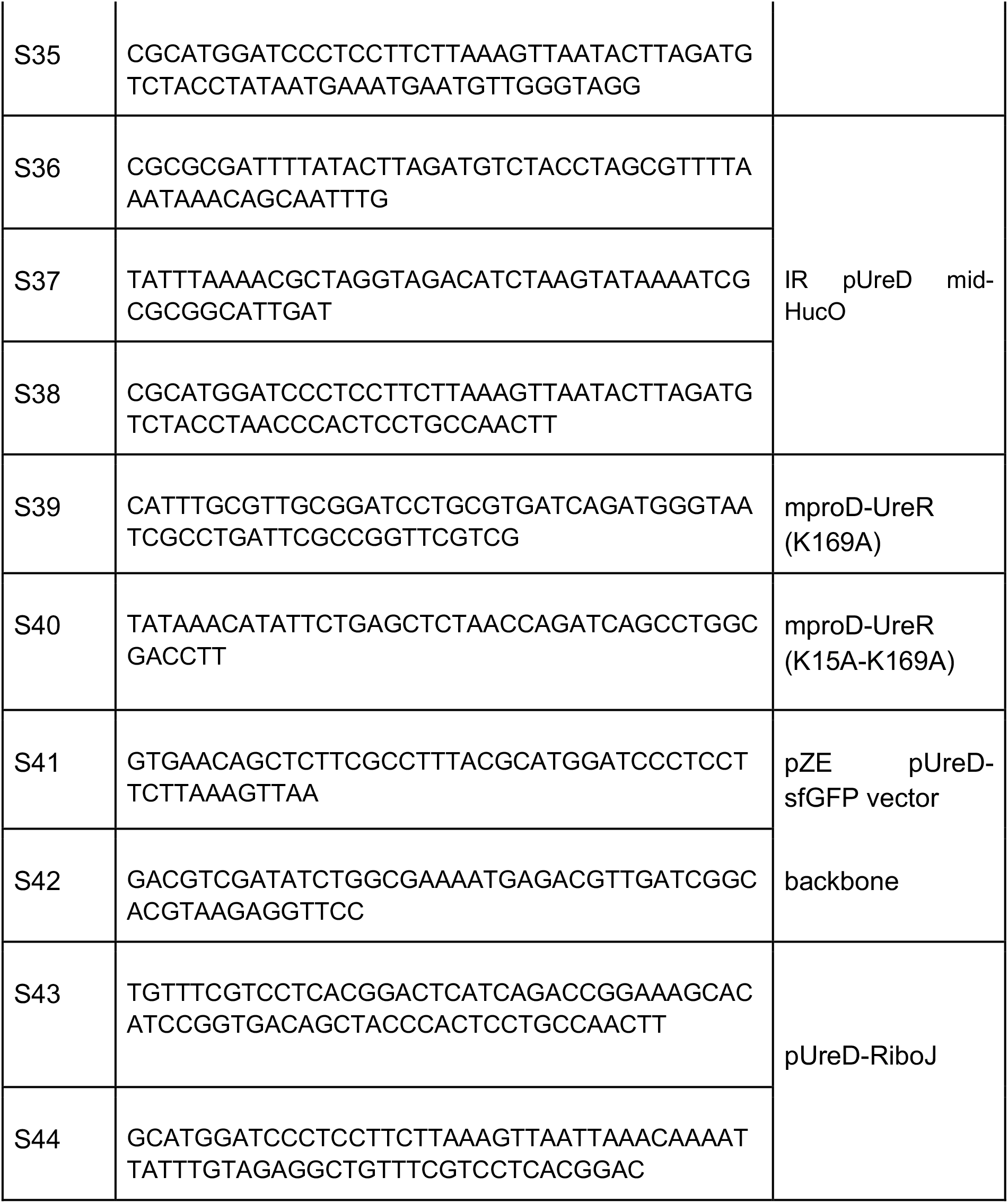
Genes and Primers used in this study.

